# The mode of expression divergence in *Drosophila* fat body is infection-specific

**DOI:** 10.1101/2020.07.30.229641

**Authors:** Bryan A. Ramirez-Corona, Stephanie Fruth, Oluchi Ofoegbu, Zeba Wunderlich

## Abstract

Transcription is controlled by the interactions of *cis*-acting DNA elements with diffusible *trans-*acting factors. Changes in *cis* or *trans* factors can drive expression divergence within and between species, and the relative prevalence of each can reveal the evolutionary history and pressures that drive expression variation. Previous work delineating the mode of expression divergence in animals has largely used whole body expression measurements in a single condition. Since *cis*-acting elements often drive expression in a subset of cell types or conditions, these measurements may not capture the complete contribution of *cis*-acting changes. Here, we quantify the mode of expression divergence in the *Drosophila* fat body, the primary immune organ, in several conditions. We performed allele-specific expression analysis using two geographically distinct lines of *D. melanogaster* and their F1 hybrids. We measured expression in the absence of infection and in separate infections with Gram-negative *S. marcescens* or Gram-positive *E. faecalis* bacteria, which trigger the two primary signaling pathways in the *Drosophila* innate immune response. The mode of expression divergence strongly depends on the condition, with *trans-*acting effects dominating in response to Gram-positive infection and *cis*-acting effects dominating in Gram-negative and pre-infection conditions. Expression divergence in several receptor proteins may underlie the infection-specific *trans* effects. Before infection, when the fat body has a metabolic role, there are many compensatory effects, changes in *cis* and *trans* that counteract each other to maintain expression levels. This work demonstrates that within a single tissue, the mode of expression divergence varies between conditions and suggests that these differences reflect the diverse evolutionary histories of host-pathogen interactions.

## Introduction

Differences in gene expression are believed to be major drivers of phenotypic divergence in closely related species (King and Wilson 1975). These differences can arise through sequence changes in *cis-*regulatory elements, such as enhancers, or in the coding regions of *trans*-acting factors, such as transcription factors. Evolutionary processes rely on both changes in *cis* and changes in *trans*, and the prevalence and relative contributions of *cis* and *trans* changes are actively being explored in various model systems (Signor and Nuzhdin 2018). For example, within individual *Drosophila melanogaster* lines or between *Drosophila* species, the contributions of *cis-*acting changes generally increase with phylogenetic distance, and the precise balance of *cis* versus *trans* effects depend on the phylogenetic relationships and demographics of the genotypes being compared (Wittkopp et al., 2004, Wittkopp et al., 2008, McManus et al., 2010, Coolon et al., 2014, Osada et al., 2017). These studies have elucidated mode and tempo of the evolutionary processes driving gene expression divergence; however, most of these studies use whole body measurements, thus averaging signal across multiple tissue and cell types. Therefore, these studies cannot examine the prevalence of *cis* and *trans* changes in specific biological processes, which may be subject to different types of selection pressure. In addition, given that many *cis-*regulatory elements act in a tissue-specific manner, studies that measure *cis* and *trans* effects with tissue-specific resolution may reveal effects that are undetectable in heterogenous samples.

*Drosophila* have an innate, but not adaptative, immune response, and this response is a powerful system for measuring the contributions of *cis* and *trans* changes for several reasons. First, the immune response is inducible, with active and inactive states. This allows for the clear delineation of the transcriptional response of the immune system from that of other processes. Second, the fat body within the immune system is an optimal tissue for study. Though other tissues participate in the immune system, the fat body is a primary driver of the humoral response (Buchon et al., 2014) and is relatively easy to isolate. Lastly, there is ample variation in the resistance, survival, and transcriptional response to infection between individual *D. melanogaster* lines (Lazzaro et al., 2004, Lazzaro et al., 2006, Sackton et al., 2010, Hotson and Schneider 2015), suggesting there are many sequence changes driving these differences.

To quantify changes in *cis* and *trans* that drive transcriptional divergence in the immune response, we used allele-specific expression analysis (ASE) of RNA-seq data (Wittkopp et al. 2004, Signor and Nuzhdin 2018, Frochaux et al 2020). In this approach, we compare a gene’s expression levels in two parental lines to the expression levels of each parental allele in the resulting F1 hybrids. Differences in expression due to changes in *cis*, for example a sequence change in a promoter or enhancer, will only affect the expression of the corresponding parental allele. Thus, changes in *cis* are independent of cellular environment and will be observed as allelic imbalance between the parents that is then maintained in the hybrids. Differences in *trans*, for example a coding sequence change in a transcription factor, will affect the expression of both alleles in the F1 hybrids and thus will be observed as differential expression in the parental lines that is not maintained in the F1 hybrids. Combining ASE with RNA-seq allows us to determine the prevalence of *cis* and *trans* changes genome-wide.

When comparing the innate immune response of different *D. melanogaster* lines, it is not clear whether *cis* or *trans* changes will dominate. Changes in *cis* generally affect a single gene’s expression and thus may easily tolerated, as they only introduce small amounts of phenotypic variation into a system. Changes in *trans* can affect the expression of many genes at once and thus efficiently introduce a large amount of phenotypic variation but may be harder for the organism to tolerate. However, the specific biology of the innate immune response may temper this expectation. Antimicrobial peptides (AMPs) are among the most highly up-regulated genes in response to infection; however, changes to their expression, and even deletion of individual AMP genes, often have little to no measurable effect of infection survival (Hanson et al., 2019). This suggests that to get an appreciable phenotypic effect, synchronous changes in gene expression are required, which can result from a single change in *trans*-acting factors. In addition, previous work has suggested that within *Drosophila melanogaster* lines, *trans* changes are typically more prevalent (Wittkopp et al., 2008, Coolon et al., 2014).

To measure the contributions of *cis* and *trans*-acting changes in the *Drosophila* innate immune response, we measured fat body gene expression in two sequenced inbred *Drosophila* lines and their F1 hybrids in control and infection conditions. We separately infected the animals with either Gram-positive *Enterococcus faecalis* or the Gram-negative *Serratia marcescens* to trigger the two primary immune signaling pathways in the fly. Using ASE analysis, we quantified the contribution of *cis* and *trans* effects in the control and in each infection condition and found that *cis* effects dominated the expression divergence among control and *Enterococcus faecalis-*infected samples, while *trans* effects were dominant in the *Serratia marcescens*-infected samples. Further analysis suggested that expression differences in several receptor proteins may be driving the observed *trans*-acting changes. In sum, our work suggests that the relative importance of *cis* and *trans* acting changes may be highly dependent on the dominant biological process, even within a single tissue.

## Results

### Two geographically-distinct lines show ample genotype-specific immune response

To measure the relative contributions of *cis-* and *trans*-acting effects in the innate immune response, we needed two inbred, sequenced strains of *D. melanogaster* with abundant genetic variation and phenotypic differences in the immune response. The founder lines of the *Drosophila* Synthetic Population Resource are inbred, sequenced, and genetically diverse, making them ideal candidates (King et al. 2012). To maximize the likelihood of finding both genetic and phenotypic variation in these lines, we selected two lines from different continents, the A4 line, also known as KSA2, collected from the Koriba Dam in South Africa, and the B6 line, collected from Ica, Peru. Using the available SNP data, we found 462,548 SNPs between A4 and B6, with about half of them falling into exonic regions, indicating that 279,656/30,107,083 = 0.9% of exonic bases varied between the genotypes. The extensive variation in the coding regions allowed us to map, on average, 11.2% (±1.3%) of RNA-seq reads in an allele-specific manner.

To assess the divergence in the A4 and B6 immune responses, we measured gene expression pre- and post-infection in the abdominal fat body, the primary site of immune response. To do so, we performed RNA-seq on the dissected fat bodies of 4-day old males from both lines that had been infected with either Gram-positive *Enterococcus faecalis* (*Efae*) or Gram-negative *Serratia marcescens* (*Smar*). We selected these bacteria because in *D. melanogaster*, Gram-positive infections generally stimulate the Toll pathway, and Gram-negative infections generally stimulate the IMD pathway, though there is additional nuance due to signaling crosstalk and the contributions of other signaling pathways (Buchon et al., 2014; Busse et al., 2007; Lemaitre and Hoffmann 2007; Tanji et al., 2010; Troha et al., 2018). We measured expression pre-infection and 3 hours post-infection, to capture the early transcriptional response prior to the complicating effects of feedback.

In response to *Efae* infection, we found sizable genotype-specific effects in the immune response. To detect these effects, we performed two types of differential gene expression analysis: we first compared control and infected samples to find *Efae*-responsive genes, and then within this group, we looked for genes differentially expressed between the A4 and B6 genotypes. We found 1165 differentially expressed genes between the control and infected samples regardless of genotype (Figure 1A). We categorized these *Efae*-responsive genes into three groups based on their differential expression between genotypes. Group 1 genes are differentially expressed only in the control samples, Group 2 genes are differentially expressed only in the infected samples, and Group 3 genes are differentially expressed in both control and infected samples. Genes not categorized into any of these groups were designated as not showing genotype-specific expression. Of the 500 *Efae*-responsive genes showing genotype effects, 87% (433 genes) are in Group 2, while only 10 genes are in Group 1 and 57 genes in Group 3 (Figure 1B). This indicates that many *Efae*-responsive genes show genotype-specific expression, and these differences are typically only revealed in response to infection.

**Figure 1.**
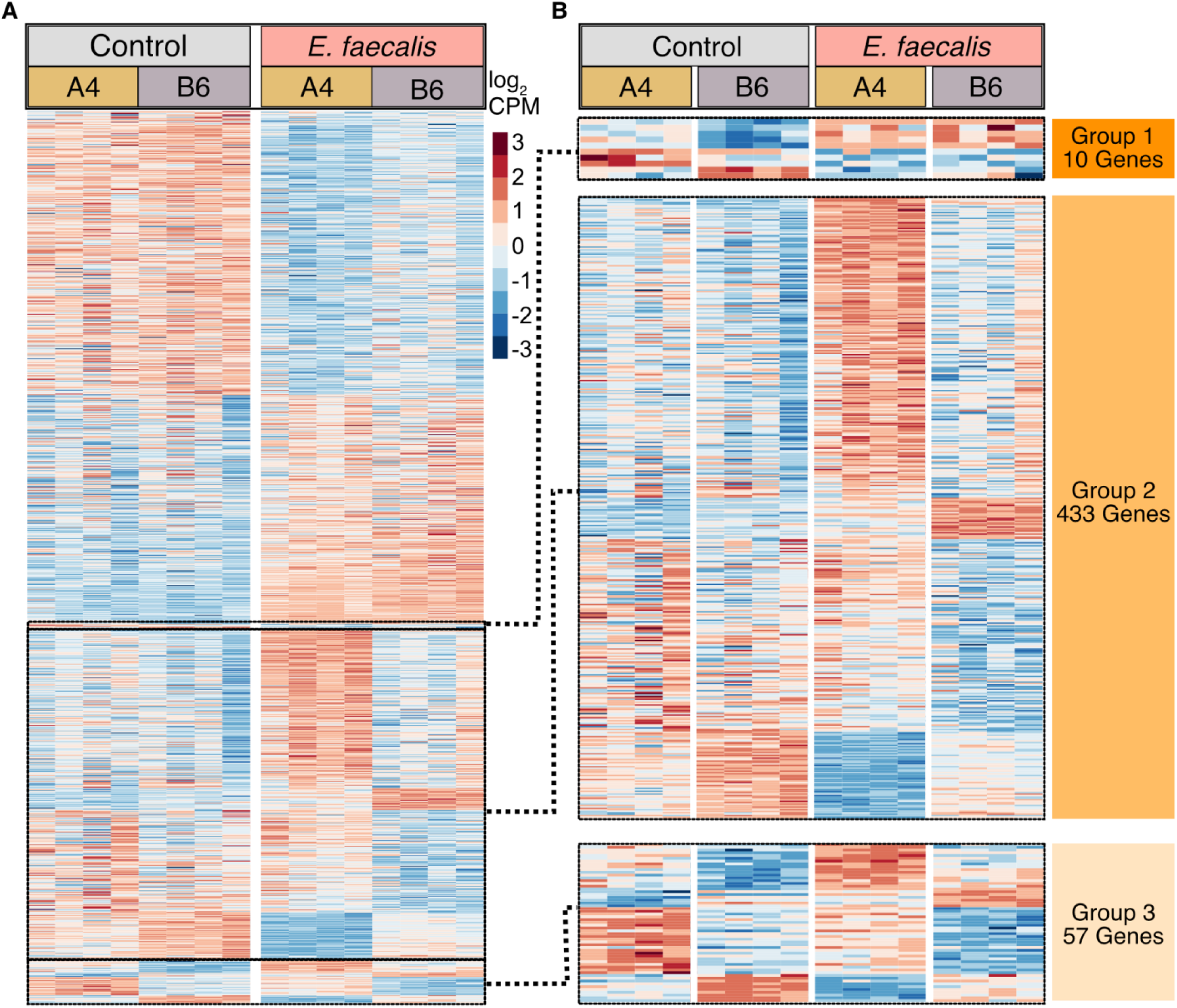
The A4 and B6 *D. melanogaster* lines have variation in their response to Gram-positive *E. faecalis* infection. A) We measured expression in the fat bodies of the A4 and B6 lines infected with Gram-positive *Enterococcus faecalis* 3 hours post-infection. Of 11038 genes detected, we found 1165 differentially expressed in response to infection, relative to control samples, of which 201 were previously published *Drosophila* immune genes. Mean centered log_2_ CPM values are displayed. B) We categorized the 1165 *Efae-*responsive genes into three groups, based on their differential expression between the two fly genotypes: genes showing genotype-specific expression only in the control condition (Group 1), genes showing genotype-specific expression only in the infected condition (Group 2) and genes showing genotype-specific expression in both control and infected conditions (Group 3). The majority of genes fell into the Group 2 classification, indicating a large amount of genotype-specific expression variation is revealed upon infection with *Efae*. The fewest genes were classified into Group 1, suggesting that in the resting state there are few genotype-specific differences in the *Efae*-responsive gene set.

In response to the *Smar* infection, we found 1203 differentially expressed genes between the control and infected samples (Figure 2A). To look for genotype-specific expression, we categorized the 1203 *Smar*-responsive genes into the three previously mentioned groups. For this infection, we found roughly equal numbers of genes in each group, with 89, 91, and 84 genes in Groups 1-3, respectively (Figure 2B). This indicates that a higher fraction of *Smar-*responsive genes show genotype effects prior to infection than *Efae-*responsive genes (p = 1.1e-11, Chi-square test, Bonferroni corrected), while a higher fraction of *Efae-*responsive genes show genotype effects after infection (p = 9.5e-67, Chi-square test, Bonferroni corrected)

**Figure 2.**
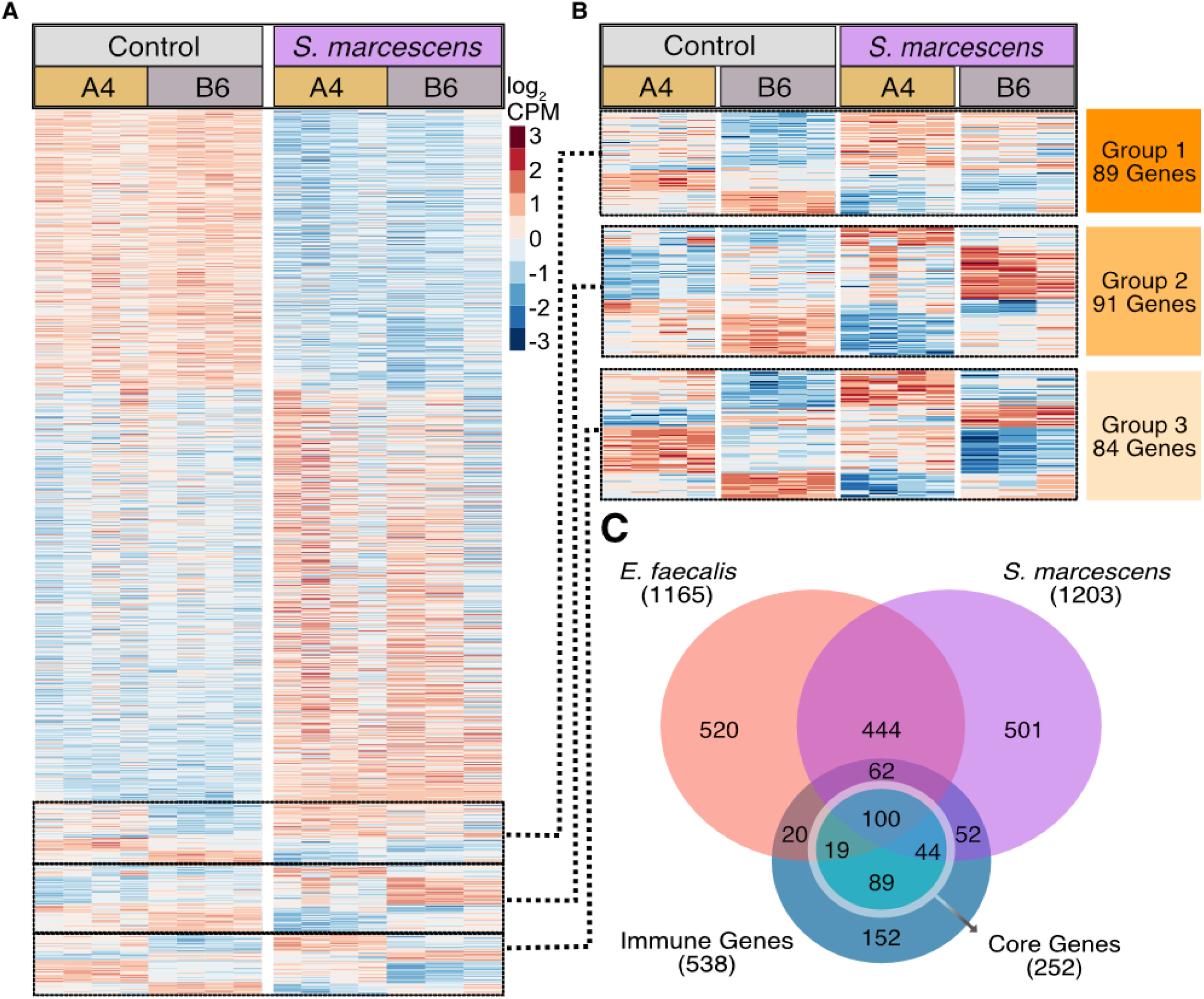
The A4 and B6 *D. melanogaster* lines vary in their response to Gram-negative *S. marcescens* infection. A) We measured expression in the fat bodies of the A4 and B6 lines infected with Gram-negative *Serratia marcescens*. Of 11071 genes detected, we found 1205 differentially expressed genes between the control and infected samples; 258 were previously described *Drosophila* immune genes. Mean centered log_2_ CPM values are displayed. B) We categorized the 1205 genes into the three previously described groups (Figure 1). Among *Smar*-responsive genes, roughly equal numbers show expression differences between the genotypes before (Group 1), after (Group 2), and both before and after infection (Group 3). C) We intersected the genes we identified as differentially expressed in response to infection and a list of previously published immune responsive genes. This list of immune genes is an expanded version of the *Drosophila* Immune Responsive Genes Set (DIRGS). More than half of the verified immune genes were identified as differentially expressed in the abdominal fat body, with half of these immune genes being shared between conditions. Among these previously identified immune genes, core genes are differentially expressed across all infections. We detected roughly 40% of the core set as differentially expressed in both our infection conditions, despite differences in the genetic background, tissue type, and time point used in our study versus previous work.

To assess whether there is also phenotypic divergence on the organismal level, we performed the *Efae* and *Smar* infections and measured survival and bacterial load for 7 days post-infection. In response to *Efae* infection, we found differences in the ability to survive infection between genotypes, with B6 showing greater ability to survive infection over A4 (Supplemental Figure S1A). In response to *Smar*, we found that while there were no significant differences in survival, bacterial load was lower in A4 than in B6 (Supplemental Figure S1B, S1C). Together, these data demonstrate that there are differences between the two lines in their ability to resist or survive infection, and that these differences are pathogen specific.

To compare our tissue-specific measurements to previous work, we intersected our *Efae-* and *Smar-*responsive genes to an existing list of immune-responsive genes. This list is an expanded version of the *Drosophila* immune responsive genes set (DIRGS) and constitutes the summation of more than two decades of work in *Drosophila* (De Gregorio et al., 2001; Lemaitre and Hoffman 2007; Troha et al., 2018). Of 538 genes on this expanded list, we found more than half of these (297 genes) were identified as immune-responsive in our data (Figure 2C). Troha and colleagues identified a subset of immune-responsive genes as the core of the immune response, i.e. the set of genes that is differentially expressed regardless of the type of bacterial infection (Troha et al., 2018). We found that of these 252 core genes, approximately 40% were found to be both *Smar-* and *Efae-*responsive in our data. CrebA, which was identified as a core gene essential for both Toll and IMD-drive immune response, is one of the genes found in this overlap. Therefore, despite differences in the genetic background, tissue (previous studies were typically done with whole body sampling), and time points, our findings show concordance with previous studies of gene expression in response to infection. More importantly, we show that the A4 and B6 lines have phenotypic divergence both in expression of immune-responsive genes and the ability to fight infection, making them suitable for subsequent F1 hybrid experiments.

### *Cis*-acting effects dominate expression variation in the uninfected fat body

To effectively quantify *cis* and *trans* effects, we needed to verify our ability to accurately analyze the allelic expression in F1 hybrids. To do so, we used the RNA-seq data from the A4 and B6 parental lines mentioned above as well as data from the F1 hybrids (A4♂x B6☿) and reciprocal crosses (B6♂x A4☿), in the control, *Efae*-infected, and *Smar*-infected conditions. Since we are using males, if our allele-specific expression analysis is correct, none of X-chromosome reads should map to the paternal genotype. Using the published A4 and B6 genomes and the Allele-Specific Alignment Pipeline (ASAP) (Krueger, https://www.bioinformatics.babraham.ac.uk/projects/ASAP/), we quantified the fraction of X-chromosome reads that incorrectly map to the paternal genotype. On average, samples had 0.5% mis-assigned reads (standard deviation = 3%), with the highest fraction being 1.2% (Supplemental Table S1). The consistent, low level of mis-assigned reads verifies our ability to accurately quantify allelic expression.

We next sought to quantify *cis* and *trans* effects in the control samples. We used the complete set of parental RNA-seq reads and the subset of the F1 hybrid reads that could be assigned to a specific allele. Using three separate generalized linear models, we tested for differential expression in the parents, allelic imbalance in the F1 hybrids, and *trans* effects between parents and F1 hybrids (see Methods) (Davidson and Balakrishnan, 2016; Osada et al., 2017; Takada et al., 2017). We then categorized each gene into one of six categories (Figure 3A). Genes showing no differential expression in the parents or F1 hybrids have no evidence of *cis* or *trans* effects and are called ***conserved***. Genes showing differential expression in both the parents and F1 hybrids and no *trans* signal are categorized as ***cis-only***. Genes showing differential expression in the parents and not the F1 hybrids are categorized as ***trans-only***. Some genes show evidence of both *cis* and *trans* effects and are categorized as either ***compensatory*** (if the changes have opposing effects on expression) or ***cis* + *trans*** (if the changes are coherent). Genes that do not fall into any of these categories have an ambiguous pattern of divergence and are called ***undetermined***.

**Figure 3.**
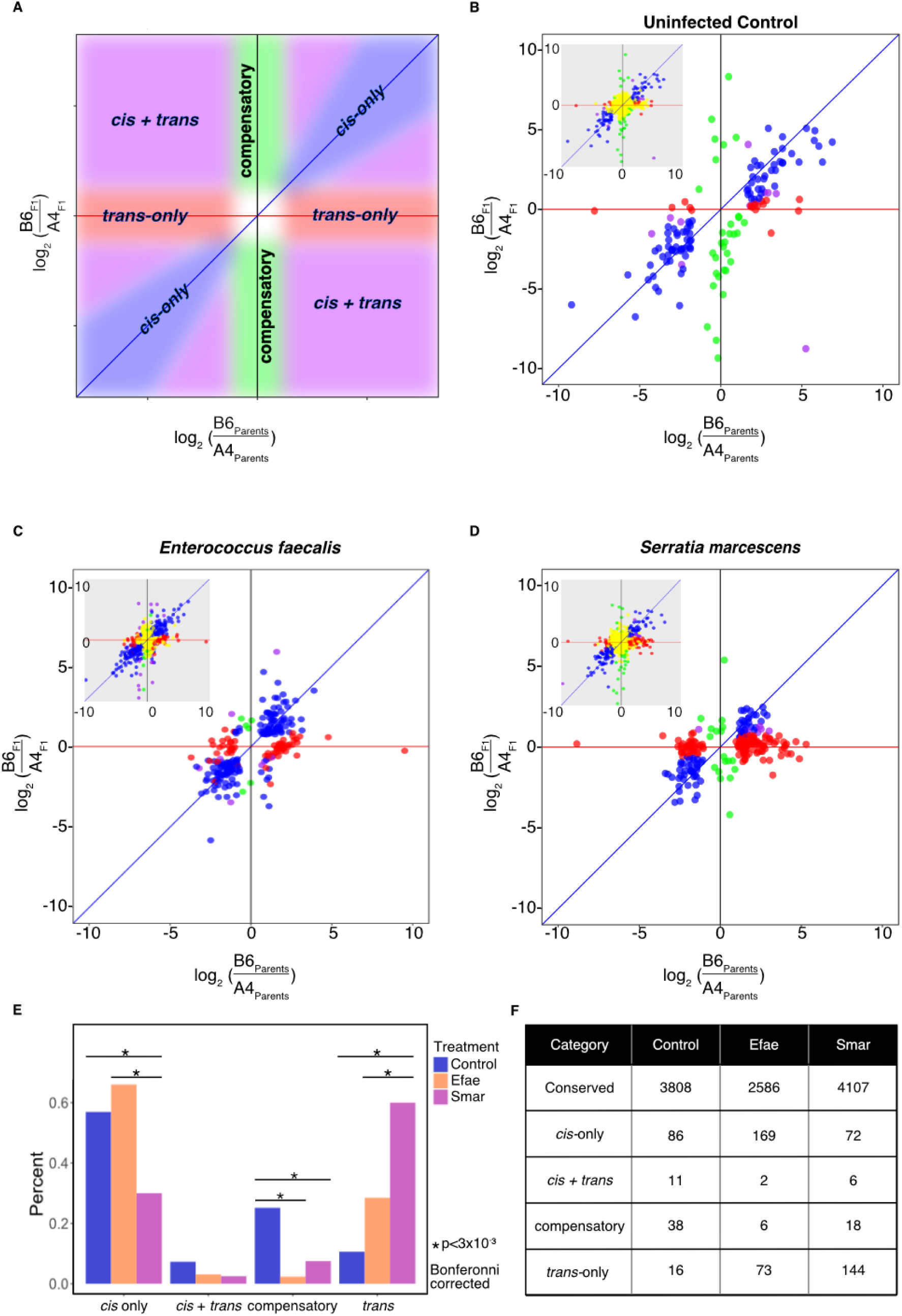
The relative contributions of *cis* and *trans* effects to expression divergence are condition specific. A) Here we show a schematic of the expected locations of four of the six categories of causes of expression divergence, conserved (yellow) and undetermined genes are excluded from downstream analysis. B) In the uninfected control condition, of 4960 genes that could be detected, 151 genes showed *cis* or *trans* signal. 86 genes show *cis*-only effects, 16 genes show *trans*-only effects, 11 genes show *cis* and *trans* effects, and 38 show compensatory effects. C) In response to *Efae* infection, expression divergence is driven predominantly by changes in *cis*. There are 256 genes that show *cis* or *trans* signal and do not overlap with the 151 genes showing signal in the control condition. Of 256 genes, 169 genes show *cis-only* signal, 86 genes show *trans-only* signal, 8 genes show a combination of *cis* and *trans* effects, and 6 genes show compensatory effects. D) In response to *Smar* infection, expression divergence is dominated by changes in *trans*. There are 240 genes that show *cis* or *trans* signal not found in the uninfected control. In these 240 genes, 72 genes showed *cis*-only signal, 144 genes showed *trans*-only signal, 6 genes showed both *cis* and *trans* effects, and 18 genes show compensatory effects. E) We compared the fraction of genes categorized into each divergence class in the three conditions and found that the modes of expression divergence were condition-specific.

Of the 4960 genes that were both expressed in the pre-infection fat body and could be detected in an allele-specific manner, 77% were categorized as conserved (3808 genes; Figure 3B). We found 151 genes showing unambiguous *cis* or *trans* effects. In these 151 genes, *cis* effects dominated the signal: 90% of genes (135 genes) showed *cis* signal (including *cis*-*only, cis + trans* and compensatory genes), and 57% (86 genes) showed *cis-only* effects. 43% of genes (65 genes) showed *trans* signal and only 10% of genes (16 genes) showed *trans-only* effects. One-quarter of genes (38 genes) were categorized as compensatory, even when using non-overlapping samples to detect *cis* and *trans* effects, which avoids the artificial inflation of compensatory signal (Fraser et al., 2019; Zhang and Emerson, 2019). Additionally, to ensure that any differences in the quality of our in-house A4 and B6 transcriptomes do not affect our conclusions, we quantified *cis* and *trans* effects using sets of high confidence genes at multiple levels of stringency and found that this had negligible effects on the detected signal (Methods; Supplemental Figures S2; Supplemental Table S2). From these data, we can conclude that in the unstimulated state, most genes have conserved expression levels in the fat body, and among those genes that diverge, *cis* effects dominate, with a sizable number of genes showing compensatory *cis* and *trans* changes.

### More *cis* than *trans* effects are found in *Efae*-infected fat body expression

We quantified *cis* and *trans* effects in *Efae*-infected samples following the same methodology. We found roughly 52% of genes (2586 genes) showed no evidence of *cis* or *trans* effects and 381 genes showed unambiguous *cis* or *trans* effects (Figure 3C). To identify genes whose expression divergence is specific to the immune response, we eliminated genes that show *cis* or *trans* signal in the control sample. After this filtering, roughly 88% of the genes showing cis or trans effects (336 genes) remained; 66% of these genes (169 genes) show *cis-only* signal, and 28% (73 genes) show *trans-only* signal. Only 8 genes (3%) show a combination of *cis* and *trans* effects, with only 6 genes showing compensatory effects. Of the genes that show *cis-only* signal, roughly even numbers of genes show higher expression in each genotype, consistent with the idea that *cis*-acting changes affect a single gene at a time. In contrast, of the genes showing *trans-only* signal, nearly twice as many were expressed more highly in the B6 genotype (47 genes) than in the A4 genotype (26 genes) (*p* = 0.00962, Chi-square test), suggesting that one or a few changes in upstream regulatory factors are responsible for this observation. Since we do not observe this trend towards higher B6 expression in the control samples and have removed genes that showed any evidence of mapping bias between the two genotypes (Methods), we are confident this trend is not an artifact and reflects true biological differences in the immune response. In sum, we find both *cis* and *trans* effects drive *Efae*-responsive expression divergence, with *cis* effects dominating and few genes showing compensatory changes.

### *Trans* effects dominate expression variation in the *Smar*-infected fat body

Lastly, we quantified *cis* and *trans* effects in response to *Smar* infection. We found roughly 82% of genes (4107 genes) are conserved, and 357 genes showed unambiguous *cis* or *trans* signal (Figure 3D). We again filtered out genes that show *cis* or *trans* effects in the control samples and were left with 240 genes that have immune-specific signal. Of these, 30% (72 genes) showed *cis-only* signal, and roughly equal numbers of *cis-only* genes showed higher expression in each genotype. 10% (24 genes) showed both *cis* and *trans* effects, and within these genes, 18 genes had compensatory signal. Surprisingly, 60% of genes (144 genes) showed *trans-only* signal. Within *trans-only* genes, we found that 71% showed greater expression in B6. Though the *Efae* analysis found far fewer genes affected in *trans*, the proportion of *trans*-*only* genes showing higher B6 expression was similar. This suggests that some of the upstream differences giving rise to the *trans* effects may be shared between the two infection conditions. In summary, in response to *Smar* infection, *trans* effects drive the majority of expression divergence between the two genotypes and few genes show compensatory effects.

### Comparisons of *cis* and *trans* signals in different conditions reveal both infection-specific and shared divergence

To systematically assess how *cis* and *trans* effects contribute to expression variance in the same tissue under different conditions, we compared the proportion of genes falling into the different divergence categories. The control and *Efae-*infected samples had a greater proportion of *cis-only* genes than the *Smar* samples (control vs. *Smar p* = 1.97e-06, *Efae* vs. *Smar p* = 1.97e-14, Chi-square test, Bonferroni-corrected). However, all three groups differ in the proportion of *trans-only* genes, with *Smar-*infected samples showing more than twice the proportion of genes with *trans*-only signal, followed by *Efae*, and then the control samples (control vs. *Efae p* = 3.7e-04, control vs. *Smar p* < 2.2e-16, *Efae* vs. *Smar p* = 2.79e-11, Chi-square test, Bonferroni-corrected). We also found that the uninfected fat body showed significantly more compensatory signal than either infected sample (control vs. *Efae p* = 2.35e-11, control vs. *Smar p* = 2.26e-05, Chi-square test, Bonferroni-corrected). Taken together, this suggests that before infection, when the fat body is carrying out its metabolic functions, there is less pressure for expression divergence, as supported by the large number of compensatory changes and conserved genes. In genes that show expression divergence in the control condition, *cis*-acting changes, which have local, non-pleiotropic effects, dominate. In response to infection, there is ample expression divergence, which is driven by both *cis* and *trans* effects. The extent to which each type of effect contributes is dependent on the particular pathogen, suggesting that the relative importance of local and pleiotropic changes is specific to different infection pressures.

Though we generally expect the two infections to regulate gene expression via distinct signaling pathways, we also anticipated some genes would be regulated in both infections, either due to crosstalk between the IMD and Toll pathways (Busse et al., 2007; Tanji et al., 2010) or via more general infection and wound responses. We found 75 genes with unambiguous *cis* and/or *trans* signal in response to both *Efae* and *Smar* infection (Supplemental Data). Of these genes, 61 showed concordant classification and the remaining 14 genes did not. Thereofre, in the majority of genes shared between these two infections, the same genetic differences are likely driving the expression divergence in both infection conditions. Additionally, in rare cases, a single gene can experience either *cis* or *trans* effects depending on the infection context.

### Differential expression of detection genes is a likely source for genotype expression bias in observed *trans* effects

Since we observed that genes with *trans*-*only* effects tended to be more highly expressed in B6 than in A4 in both infection conditions, we hypothesized that changes in a handful of upstream immune factors are responsible for this phenomenon. The changes in upstream regulators can either be infection-specific or shared, and genes affected by shared regulators would show *trans* signal in both infections. Out of 217 genes showing *trans-only* signal in either infection, only 13 genes were shared, 4 with higher expression in A4, and 9 with higher expression in B6. The small number of genes that show *trans* effects in both infection conditions indicates that the bulk of *trans*-acting changes are likely infection-specific and not driven by a shared infection or wound healing response.

To find likely sources of infection-specific *trans* effects, we hypothesized that immune detection genes, signaling genes, or transcription factors differentially expressed between genotypes in the control condition would be likely candidates, since these genes have the ability to affect the expression of many downstream targets. Further, we posited that these genotype-specific differences had to be present in the control to have the effects at the 3-hour post-infection timepoint. Of the approximately 300 genes that are differentially expressed between genotypes in the control samples, we found 22 genes that are prime candidates (Table 1). Fourteen are previously identified immune-responsive genes, and eight are transcription factors not yet implicated in immune response, but that may be peripherally involved.

**Table 1:** Transcription factors and immune genes identified as potential sources of trans effects in infection. List of genes potentially involved in observed *trans* effects for *Efae* and *Smar* infection. Candidate genes were identified by finding genes that had genotype-specific expression differences in the uninfected control conditions and were classified as either a transcription factor, immune signaling gene, or immune detection gene.

| FB Gene ID | Gene name | Type | Log<sub>2</sub> Fold Change (B6/A4) | More Highly Expressed in: | A4 Average CPM | B6 Average CPM | Immune involvement |
| --- | --- | --- | --- | --- | --- | --- | --- |
| FBgn0029822 | CG12236 | TF | -3.13 | A4 | 28 | 2.7 | Unclear |
| FBgn0039075 | CG4393 | Signaling | 2.06 | B6 | 8.8 | 39 | Unclear |
| FBgn0038978 | CG7045 | TF | 3.06 | B6 | 7 | 57 | Unclear |
| FBgn0001981 | esg | TF | -3.27 | A4 | 11 | 1 | Unclear |
| FBgn0039932 | fuss | TF | 2.50 | B6 | 1.1 | 6.3 | Unclear |
| FBgn0250732 | gfzf | TF | 8.24 | B6 | 0 | 2 | Unclear |
| FBgn0000448 | Hr46 | TF | -4.12 | A4 | 10 | 0.7 | Unclear |
| FBgn0016675 | Lectin-galC1 | Detection | 2.72 | B6 | 79 | 570 | Binding and agglutination |
| FBgn0035993 | Nf-YA | TF | -10.16 | A4 | 10 | 0 | Unclear |
| FBgn0028542 | NimB4 | Detection | -1.08 | A4 | 40 | 22 | Unclear |
| FBgn0259896 | NimC1 | Detection | -3.06 | A4 | 97 | 27 | Unclear |
| FBgn0003130 | Poxn | TF | -4.38 | A4 | 1.2 | 0.07 | Unclear |
| FBgn0014033 | Sr-CI | Detection | -2.39 | A4 | 84 | 38.6 | Unclear |
| FBgn0004606 | zfh1 | Signaling / TF | 1.77 | B6 | 50 | 17 | Unclear |
| FBgn0031973 | Spn28Dc | Signalin<br>g | 2.56 | B6 | 9.4 | 2.4 | Negative regulator of melanization |
| FBgn0037906 | PGRP-LB | Detectio<br>n | 4.536 | B6 | 111.9 | 75.2 | Negative regulator of IMD pathway |
| FBgn0043576 | PGRP-SC1a | Detectio<br>n | -5.57 | A4 | 4.3 | 6.8 | Negative regulator of IMD pathway |
| FBgn0033327 | PGRP-SC1b | Detectio<br>n | -5.24 | A4 | 3.9 | 0.2 | Negative regulator of IMD pathway |
| FBgn0043575 | PGRP-SC2 | Detectio<br>n | -4.02 | A4 | 15 | 1.3 | Negative regulator of IMD pathway |
| FBgn0035806 | PGRP-SD | Detectio<br>n | 4.25 | B6 | 97.6 | 19.2 | Positive regulator of IMD pathway |
| FBgn0039102 | SPE | Signalin<br>g | 2.41 | B6 | 491.9 | 255.2 | Positive regulator of Toll pathway |
| FBgn0003495 | spz | Signalin<br>g | 0.68 | B6 | 72.8 | 45.5 | Positive regulator of Toll pathway |

Five peptidoglycan recognition proteins (PGRP) genes are potential mediators of the large number of *trans* effects observed in the *Smar* infection. Four of these PGRPs (PGRP-SC1a, PGRP-SC1b, PGRP-SC2, PGRP-LB) are negative regulators of the IMD response, and the last gene, PGRP-SD is positive Toll and IMD regulator (Bischoff et al., 2006; Zaidman-Rémy et al., 2006; Iatsenko et al., 2016; Charroux et al., 2018; Lu et al., 2020). Three of the negative regulators, PGRP-SC1a, PGRP-SC1b, PGRP-SC2, are more highly expressed in A4. Given that these are negative regulators of the IMD pathway, this finding is congruent with the observation that genes showing *trans-only* signal tend to show greater expression in B6. PGRP-SD is more highly expressed in B6, and, given its role as a positive regulator of the IMD response, it is also consistent with the trend of higher expression of genes showing *trans-only* signal in the B6 line. The last negative regulator of IMD response, PGRP-LB, was also found to have higher expression in B6. Since there are more negative regulators showing higher expression in A4 than B6, it is possible the balance of negative to positive regulators can account for the expression trend observed in *Smar trans-only* genes. It is also possible that the greater expression of PGRP-SD is enough to account for the differences observed.

Though there were fewer *trans* effects in the *Efae*-infected samples than in the *Smar-*infected samples, the pattern wherein nearly twice as many genes showed greater expression in B6 than A4 was maintained. Of the 22 candidate genes, we found two Toll-specific genes that may be responsible for the observed signal: *Spatzle-Processing Enzyme* (*SPE*) and *spatzle* (*spz*), which are more highly expressed in B6. *Spatzle* is the Toll receptor ligand, and *SPE* is required to generate the active form of *spz*, so differential expression of these genes can drive a large number of downstream changes. In addition, PGRP-SD can act as a positive regulator of both the Toll and IMD responses and is also found to have higher expression in the B6 line.

In addition to differences in expression levels between genotypes, function-altering differences in the coding sequences of key immune genes may also be the source of the observed *trans-*acting changes. To analyze the pattern of coding sequence variants, we used the Ensembl Variant Effect Predictor (VEP) to identify the proportions of synonymous to nonsynonymous coding changes between A4 and B6 in several gene sets (McLaren et al., 2016). We considered all genes expressed in the fat body above a threshold (CPM>1), and then sorted them into two groups: genes that are differentially expressed in response to either or both infections (*DE infection*) and those that are not (*fat body detected*). We also generated a gene set that is the intersection between *DE infection* and our list of curated immune-responsive genes (*DE immune*; Figure 4A). We posited that, given the large number of *trans* effects in response to infection, immune-related genes may have a greater number of nonsynonymous to synonymous changes, compared to the *fat body detected* gene set. We found that *DE immune* genes have a significantly higher fraction of nonsynonymous sequence changes (24%) compared to the *fat body detected* genes (21%) (*p* = 0.007, Chi-square test, Bonferroni-corrected), suggesting that some of these changes may be under selection and possibly the source of our *trans*-acting signal (Figure 4B-C). *DE infected* genes showed lower fractions of nonsynonymous changes (20%) compared to the *fat body detected* genes (*p* = 0.04, Chi-square test, Bonferroni-corrected). We may not see an elevated fraction of nonsynonymous changes in the *DE infected* gene set because this includes both immune-related genes, but also genes with unrelated functions whose expression may be regulated due to the metabolic constraints on the fat body tissue.

**Figure 4:**
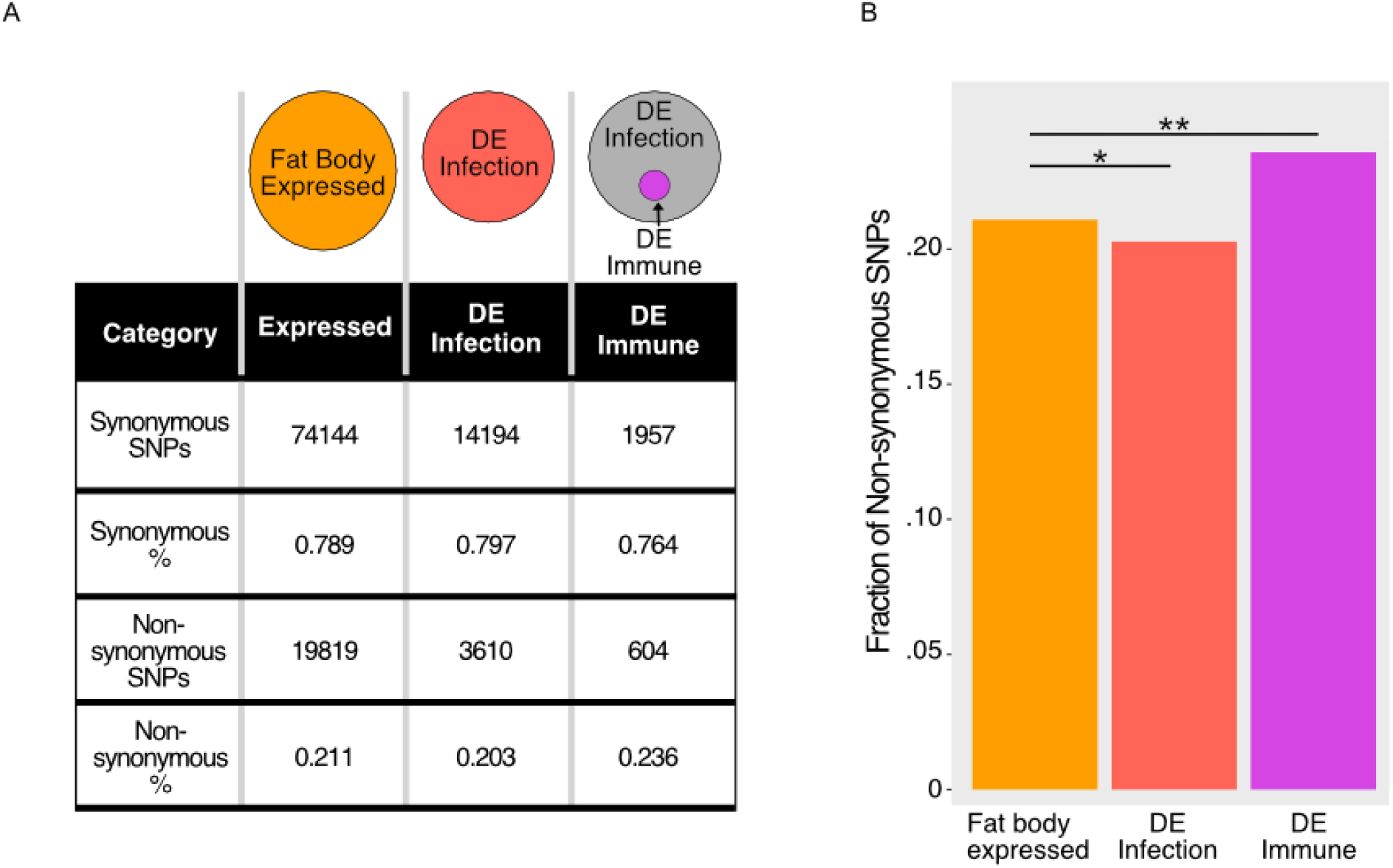
There is a greater proportion of non-synonymous SNPs in previously identified immune-responsive genes. A) To look for the prevalence of non-synonymous SNPs, we defined three gene sets. Among genes detected in the fat body samples, we separated genes into those that were differentially expressed in response to either infection (DE infection), and those that were not (fat body expressed). Among the DE infection genes, we further refined the gene list to include only previously identified immune genes (DE immune). The numbers indicate the total number of SNPs found in each gene set. B) DE immune genes have a higher proportion of non-synonymous SNPs than the fat body expressed genes, which suggests they may be subject to different selection pressures. *p*-values are Bonferroni-corrected Chi-square test with the proportion of non-synonymous SNPS relative to the fat body expressed gene set.

This analysis focuses on overall patterns of coding sequence changes in these large gene sets. We recognize that even an individual coding sequence change may drive many downstream expression differences and analyzed these changes, but no obvious candidates emerged (Supplemental Figures S3, Supplemental Table S3 and S4). Predicting the effect of these mutations on individual protein function, however, remains a challenge.

## Discussion

Here, we quantified the mode and extent of expression divergence in the *Drosophila* abdominal fat body, both in an uninfected control condition, where it carries out a variety of metabolic roles, and in response to two types of infection. We found that two geographically isolated lines of *D. melanogaster* are phenotypically distinct in their immune responses, differing both on the organismal and transcriptional levels. By comparing gene expression in the fat body between these lines and their F1 hybrids, we quantified the contributions of *cis* and *trans* effects to expression divergence in the uninfected control, *Efae-*infected and *Smar-*infected conditions. Both the control and *Efae* infection conditions were dominated by *cis* effects, while the *Smar* infection condition had an abundance of *trans* effects. Notably, the uninfected control also showed a greater proportion of compensatory effects, suggesting that there is stabilizing selection to maintain fat body expression levels of certain genes in the absence of an infection. Among the genes showing changes in *trans*, we found that expression of the B6 allele is typically higher in both types of infections. This suggests that changes to a small number of upstream immune response regulators may be responsible for this bias, and we identified expression divergence in a group of receptor programs that may drive these *trans* effects. Overall, we find that the mode of evolution in expression divergence can vary between conditions in a single tissue and likely represents condition-specific selection pressures.

Our unique approach to measuring the mode of expression divergence gave rise to several novel observations about the relative contributions of *cis* and *trans* effects on expression variation. While there have been a number of studies aimed at disentangling the contribution of *cis* and *trans* changes to gene expression in *Drosophila*, few have sought to answer this question using a single organ or with different physiological stimuli (Wittkopp et al., 2004, Wittkopp et al.,2008, McManus et al., 2010, Coolon et al., 2014, Osada et al., 2017). Our approach allows us to examine evolutionary changes in response to perturbation while minimizing the confounding effects of multiple tissue types. There are three studies most closely related to ours. A previous study by Juneja, et al. (2016) found, among geographically distinct flies, a large number of *cis*-acting changes that cause whole body expression divergence in response to an infection with mixture of bacteria. This is concordant with our finding of a large number of *cis*-acting changes in both infection conditions, but this study did not quantify *trans*-acting changes or distinguish between Toll- and IMD-specific responses. By measuring expression in the heads and abdomens of multiple *D. melanogaster* lines, another group reported the predominance of changes in *cis* over those in *trans* but did not measure these differences in different physiological states or attempt to dissect individual tissues in the head or abdomen (Osada et al., 2017). Most recently, Frochaux et al., (2020) sought to uncover the underlying genetics of *P. entomophila* resistance in the gut and identified a novel driver of this phenotype but limited their analysis to locally acting eQTLs. Here, we sought to directly assess the contribution of *cis* and *trans* sequence changes in a single tissue in the context of multiple treatment conditions, giving a uniquely high-resolution view of the evolutionary sequence changes underlying expression divergence.

With our approach we were able to uncover two notable trends. First, we found that compensatory mutations were more frequent in the control samples. We observe that while overall compensatory effects are less common than *cis-only* and •*trans-only* effects, compensatory effects in the uninfected samples are more pervasive than in either of the infected conditions. Previous studies in several organisms had suggested that compensatory effects were very prevalent (McManus et al., 2010, Gonclaves et al., 2012, Schaefke et al., 2013, Coolon et al., 2014). This was perplexing because it would seem to suggest that, even in between species, selective forces to alter expression and to stabilize it were at odds and did not explain how biological systems are able to evolve divergent expression. However, certain choices in experimental design can inflate estimates of compensatory effects (Zhang and Emerson 2019; Fraser et al., 2019). To avoid this artifact, we use non-overlapping F1-hybrid samples and therefore have generated more accurate estimates of compensatory effects across multiple conditions. Additionally, a large proportion of studies addressing *cis* and *trans* effects in animals do so in “control” conditions, which may not reveal the full extent of selection forces that act on gene expression (Gonclaves et al., 2012, Osada et al., 2017, Davidson and Balakrishnan 2016, Signor and Nuzhdin 2018). In our system, we find evidence that the genes involved in the maintenance of basic metabolic functions of the uninfected fat body are under different selective pressures than those involved in immune response. Unlike the immune-responsive genes, which must contend with a continuously evolving pathogen landscape, the genes carrying out metabolic functions may be subject to stabilizing selection, given relatively unchanging nutritional availability. In future studies, it will be interesting to further probe which systems and conditions show enrichment for these different patterns of expression divergence.

Secondly, we observe that the relative contribution of *cis-* and *trans-*acting changes are perturbation-specific. Most notably, in response to *Efae* infection, *cis* effects dominate expression changes, while in the *Smar* infection, *trans* changes are predominant. The prevalence of either *cis* or *trans* effects can be reasonably justified in our system, but we did not anticipate that the proportion of these effects would be infection specific. Because changes in *trans* factors have pleiotropic effects, it has been suggested that changes to these factors are under more selective constraint than *cis*-acting elements, and, thus, *cis* effects can more readily introduce small-scale variation into a system (Schaefke et al ., 2013). In some cases, however, arriving at a more fit phenotype may require the coordinated alteration of expression of many genes, which may be more readily achieved by changes to *trans*-acting factors. In other cases, a coordinated change in all genes affected by perturbing a *trans-*acting factor may not yield a phenotype with a net beneficial effect, and it may be more likely that changes to individual genes offer paths to higher fitness. In our *D. melanogaster* lines, *S. marcescens* is more virulent than *E. faecalis* – a higher dose of *E. faecalis* is needed to achieve similar levels of mortality to that of *S. marcescens* (Supplemental Figure S1). It is possible that adaptation to highly virulent pathogens or rapidly evolving pathogens requires large-scale, synchronous changes to expression, whereas adaptation to less virulent pathogens is possible with smaller, localized mutations. Consequently, we suggest that the differences in the abundance of *cis* or *trans* effects may reflect the individual details different host-pathogen interactions and how that influences the genetic architecture of adaption. Experiments with a wider range of pathogens will further illuminate the relationship between the mode of expression divergence and the host-pathogen relationship.

In summary, we find that the mode of expression divergence, as represented by the proportion of *cis* and *trans* effects in a system, is condition-specific in the *Drosophila melanogaster* abdominal fat body. This specificity is likely a result of the distinct selective pressures that different host-pathogen interactions exert on the *D. melanogaster* immune system. In the course of our study, we found several candidate genes that may be the sources of the observed *trans* effects, which are most prominent in *Smar* infection. To verify these effects, we aim to over-express these candidate genes in multiple genetic backgrounds in future experiments, which are becoming more feasible with the development of new genetic tools. In the longer term, we can combine the data sets presented here with other types of functional genomics experiments to identify immune-responsive *cis-*acting elements and the sequences changes that drive *cis*-acting divergence. Taken together, these studies will provide a more comprehensive view of how regulation of expression in this rapidly changing system is wired and evolves.

## Methods

### Animal genotypes, infection, and survival analysis

The A4 and B6 *D. melanogaster* lines, SNP tables and genomic reads were received from the *Drosophila* Synthetic Population Resource (King et al., 2012). Flies were reared at 25°C on standard cornmeal/yeast media (recipe available upon request). For all RNA-seq experiments, four-day-old males were infected with approximately 15 nL of A_600_ = 0.5 OD solution of either *Enterococcus faecalis* or *Serratia marcescens* via microinjection, yielding an infection of ∼10,000 CFUs/fly (Khalil et al., 2015). Uninfected controls were placed on a carbon dioxide pad for 6 minutes to mimic the effects of anesthesia used for microinjection. Both bacteria were grown in liquid culture on a shaker at 37°C overnight and then diluted 1:1000 in fresh media in the morning. These cultures were grown until exponential phase (*S. marcescens* in Luria-Bertani broth for 4 hours, *E. faecalis* in brain-heart infusion media for 5 hours). Bacteria were then pelleted down and resuspended in PBS for OD measurement and injection. All injections took place between 3:00 and 5:00 pm to account for the impact of circadian rhythm on immune response (Scheiermann et al., 2013). For the survival analysis, we used lower doses of infection to more effectively ascertain differences in survival. We infected A4 and B6 genotypes with either 5,000 CFUs of *E. faecalis* or 1,000 CFUs of *S. marcescens*, and the survival status of the flies was recorded once per day following infection (see Supplemental Figure S1 for details). To measure bacterial load, a sample of living flies were collected once per day, and we measured load with dilution plating as in (Khalil et al., 2015). Kaplan-Meier estimates of survival were calculated using the `survival` package in R (Therneau et al., 2020), and log-rank tests and plotting were performed using the `survminer` package (Kassambara and Kosinski 2019).

To determine the number of unique SNPs between A4 and B6, we downloaded published SNP tables available through the DSPR website (King et al., 2012). We selected for SNPs not shared between lines and that also showed a reference allele frequency of < 0.05 (implying an alternative allele frequency > 0.95). We then calculated total SNP differences for exonic and non-exonic regions using exon coordinates from Flybase (dm6/iso-1: FB2019_01) (Thurmond et al., 2019).

### Preparation and sequencing of RNA-seq libraries

Three hours after infection, abdominal filets with the attached fat bodies were prepared as in (Krupp and Levine et al., 2010). Three fat bodies per sample were suspended in Trizol (Life Technologies) and stored at -80°C for later extraction. RNA was extracted from samples using Zymo Research Direct-zol RNA Extraction Kits. Library construction was completed using a modified version of the Smart-Seq2 protocol outlined in (Serra et al., 2018). Samples were then sequenced on Illumina Next-seq Platform with NextSeq 500/550 High Output Kit v2.5 to generate 43bp paired end reads. Data was imported to the UCI High Performance Computational Cluster for trimming and mapping of sequenced reads.

### Differential expression analysis

Reads were trimmed and filtered using Trimmomatic 0.35 (Bolger et al., 2014). Count and TPM data for each sample was then calculated using Salmon 0.12.0 aligner (Patro et al., 2017) using the dm6/iso-1 transcriptome. Count matrices of gene-level data were then constructed in R using the `Tximport` 1.12.3 package (Soneson et al., 2015). To find genes differentially expressed in response to each infection, compared to control, we used the `EdgeR` 3.26.5 package (Robinson et al., 2010, McCarthy et al., 2012), excluding lowly expressed genes (CPM<1) with false discovery rate corrected *p*-values (Benjamini and Yekutieli et al., 2001). Genes with an FDR < 0.05 were considered differentially expressed. To determine genotype-specific effects, among the *Efae-* or *Smar*-responsive genes, we used EdgeR to find genes differentially expressed between A4 and B6 in either the control conditions or the treated condition.

### Generation of A4 and B6 transcriptome annotations

To map RNA-seq reads in an allele-specific manner, we created two reference transcriptomes by lifting over Iso-1 genome annotations to sequenced A4 and B6 genomes. Using tools from UCSC liftOver suite, custom chain files were created by mapping homologous sequences to the A4 or B6 genome (Salinas et al., 2016). To assess the quality of our annotations and remove genes with poor annotations, genomic sequencing reads from the DSPR website where downloaded and aligned to our transcriptome files (Thurmond et al., 2019). We hypothesized that well-annotated genes would show similar coverage of genomic reads in both the A4 and B6 transcriptomes. We then filtered genes using two methods for outlier calling: a Poisson distribution-based method and a negative binomial generalized linear model (GLM) method, similar to that used for differential gene expression in RNA-seq experiments. For the Poisson method, we fit a Poisson distribution to gene count data for the A4 and B6 transcriptomes separately, using the `fitdistributionplus 1.0-14` package in R and called outlier genes using three thresholds of increasing stringency p = 0.001, 0.01 and 0.025. For the GLM-based approach, we looked for gene counts that were significantly different between the A4 and B6 transcriptomes and filtered genes using FDR thresholds of 0.01, 0.05 and 0.09. As our threshold for significance became more stringent, we filtered out an increasing number of genes (Supplemental Figure S2). Genes found not to be outliers in either the Poisson or GLM method were then combined into gene sets based on the stringency of filtering. These gene sets were then used to quantify *cis* and *trans* effects for all three conditions. We found that the stringency of filtering did not significantly impact the total number or proportions of *cis* and *trans* effects between conditions. For the allele-specific expression analysis presented in Figure 3, we used a set of genes filtered using a combination of both methods at medium stringency.

### Allele-specific expression analysis

RNA reads were assigned parental alleles using Allele Specific Alignment Pipeline (Krueger, https://www.bioinformatics.babraham.ac.uk/projects/ASAP/) using the A4 and B6 genomes and allowing for no mismatches. Non-uniquely assignable reads were discarded. Count and TPM data were then generated by aligning allelic reads to the corresponding transcriptome. Count matrices of gene-level data were then constructed in R using the `Tximport` package (Soneson et al., 2015). Accuracy of allele calling was assessed by the proportion of X chromosome reads that aligned to the maternal genotype versus the paternal genotype. Given that all the flies are male, any reads aligning to the paternal X chromosome can definitively be classified as mis-assigned.

To characterize expression divergence into *cis* and *trans* categories, differential expression was determined with unparsed parental reads and allele-specific reads from the F1 hybrids, using EdgeR and three distinct GLM structures. Genes not found in both A4 and B6 transcriptomes, lowly expressed genes (CPM<1) and X chromosome genes were excluded from the analysis. For each condition, we first tested for differential gene expression between parental samples (DE parents; Murad et al., 2019). Next we tested for allelic imbalance, taking into account parent of origin and maternal genotype effects as outlined in (Osada et al., 2017; Takada et al., 2017). For this test we used half of the F1 hybrid samples. Finally, we tested for *trans* effects using parental samples and the remaining F1 hybrid samples (J. Coolon pers. comm.). In all three tests, we assigned significance after adjusted *p*-values for multiple comparisons using the False Discovery Rate method (Benjamini and Yekutieli, 2001). Using the results from each test, we categorized each gene into one of five classes using the following logic, which is based on previous studies (Emerson and Li 2010, McManus et al., 2010):

| Category | DE parents | F1 allelic imbalance | *Trans* test |
| --- | --- | --- | --- |
| *cis* only | True | True | False |
| *trans* only | True | False | True |
| *cis* + *trans* | True | True | True |
| Compensatory | False | True | True |
| Conserved | False | False | False |

Any genes that did not fit into the patterns found above were categorized as “undetermined” and were excluded from further analysis. A complete list of genes and their categories for each condition is available in the Supplemental Data.

### Identification of sources of trans effects

To investigate potential sources of observed *trans* effects, we looked for genes differentially expressed in our uninfected samples. This included genes that are differentially expressed between A4 and B6 only uninfected samples and genes differentially expressed in response to infection and effects in the control (Groups 1 and 3 from Figures 1-2). These genes were then intersected with a list of known *Drosophila* transcription factors as well as known immune genes (De Gregorio et al., 2001; Lemaitre and Hoffman 2007; Celniker et al., 2013, Troha et al., 2018). Only genes that were transcription factors, immune detection genes, or immune signaling genes were considered to be candidates.

### Analysis of SNPs

To better understand the effects of sequence changes on coding regions between our lines we used the Ensemble Variant Effect Predictor Tool (VEP) to predict the effects of SNPs on the resulting amino acid sequence (McLauren et al., 2016). We first created two mutually exclusive lists of genes. The first list consists of genes found to be expressed in the unstimulated fat body above a CPM of 1 and excluding genes found to be differentially expressed in response to infection, this list was referred to as *fat body expressed* genes. The second list consists of only genes differentially expressed in response to infection with either *Efae* or *Smar*, referred to as *DE infection*. We then subsetted the *DE infection* genes to make a list of genes that are differentially expressed genes that are also previously verified immune response genes, which we called *DE immune*. For each list of genes, we pulled out SNPs falling into coding regions and ran these through VEP. The proportion of synonymous to non-synonymous SNPs was then compared between conditions.

### Description of statistical tests

*p*-values for all single and multiple proportion comparisons were calculated using R’s prop.test function which performs a Chi-square test with Yate’s continuity correction. For data where more than one test was performed, *p*-values were Bonferroni corrected by multiplying the p-value by the number of tests performed.

## Data Access

All raw and processed sequencing data generated in this study have been submitted to the NCBI Gene Expression Omnibus (GEO; https://www.ncbi.nlm.nih.gov/geo/) under accession number GSE155033.

## Acknowledgements

We would like to thank J.J. Emerson and Xinwin Zhang for their thoughtful comments and suggestions on this work, Ali Mortazavi and Lorrayne Serra for their insight and access to sequencing equipment, Tom Schilling for access to microinjection equipment, Joseph Coolon, Carl de Boer and Rabi Murad for sharing their scripts as well as general guidance with computational protocols, and Anthony Long and Mahul Chakraborty for their insight helpful comments on the genomes utilized. This work was funded in part by the National Science Foundation, Award 1953324 to Z.W. B.R. was supported by an NSF Bridge to the Doctorate Fellowship. S.F. was supported by a UCI UROP award. O.O. was supported by NIH Grant R25 GM055246, T34 GM136498, and an UCI UROP award.

## Disclosure Declarations

The authors have no conflicts of interest to declare.

